# Marine fishes exhibit exceptional variation in biofluorescent emission spectra

**DOI:** 10.1101/2024.12.18.629237

**Authors:** Emily M. Carr, Mason M. Thurman, Rene P. Martin, Tate S. Sparks, John S. Sparks

## Abstract

Biofluorescence is a phylogenetically widespread phenomenon among marine fishes, yet the phenotypic diversity in fluorescent emission wavelengths (e.g., green, red) remains poorly studied across the broad diversity of marine teleosts. In this study we investigate the fluorescent emission spectra from a diverse array of 18 teleost families and record fluorescent emission peaks over multiple body regions. Our results show that fluorescent emission spectra are remarkably diverse among teleost families, as well as within genera. Fluorescent emissions also varied across different body regions within some individuals. We show that members of the families Gobiidae, Oxudercidae, and Bothidae, exhibit at least six distinct, non-overlapping fluorescent emission peaks. Nine of the 18 families examined in this study were found to have at least four distinct and non-overlapping fluorescent emission peaks. Further, we find that several families exhibit multiple discrete emission peaks for a single fluorescent color (i.e., wavelength range), including multiple distinct peaks within the green and red portions of the spectrum. This exceptional degree of phenotypic variation in fluorescent emissions highlights the potential for a diverse, even species-specific, fluorescent signaling system in certain teleost families. The interplay between different fluorescent emission wavelengths and notable variation in the distribution of fluorescence on the body could allow for a wide array of fluorescent patterns to be produced by an individual or among closely related species. Our results reveal far more diversity in both fluorescent emission wavelengths (colors) and in the distribution of fluorescent molecules on the body than had previously been reported in the literature. We characterize this novel variation in fluorescent emissions across an array of teleost families that exhibit biofluorescence and discuss the potential functional implications of this exceptional phenotypic variability.

## Introduction

Biofluorescence results from the absorption of high energy, shorter wavelength light by an organism and its reemission at longer, lower energy wavelengths [1]. Although present in numerous invertebrate and vertebrate lineages [1–7], biofluorescence is particularly phylogenetically widespread and phenotypically variable in marine fishes, where fluorescent emissions are generally reported within the red and green portions of the visible spectrum [1,8,9]. Fishes are hypothesized to use biofluorescence for intraspecific signaling, visual enhancement, and camouflage [1,10–12]. However, these potential functions require that conspecifics or predators can visualize fluorescent emissions, either as a color signal or as enhanced contrast (e.g., against a background or substrate) [1]. Most biofluorescent teleosts are cryptically patterned reef fishes [1] whose eyes are generally most sensitive to shorter wavelengths of blue, green, and yellow [13,14]. These wavelengths are the most common at depth, as the attenuation of sunlight through water rapidly removes longer (orange-red) wavelengths. This creates a monochromatic blue environment of 470-480 nm by around 150 m depth in clear oceanic waters [15]. One of the potential functional benefits of fluorescence in marine environments is that it restores longer wavelengths (green-red) in these habitats where only shorter blue wavelengths can penetrate [1,11]. Interestingly, many reef fishes (e.g., Pomacentridae, Gobiidae, Labridae) possess long wavelength sensitivity (LWS) opsins in their eyes, allowing them to visualize red wavelengths [13,16,17]. For example, *Eviota pellucida* (Gobiidae), a species that emits red fluorescence, has a spectral range that extends to ∼650 nm but with a spectral sensitivity that peaks at 450-550 nm [8].

Although the function of biofluorescence remains unknown in many lineages of marine fishes, recent studies suggest that it can serve as a visual aid. Bright green fluorescence was shown to significantly increase contrast at depth in catsharks, making it easier for conspecifics to see each other in these dimly lit environments [11]. Red fluorescence in the eyes of certain reef fishes or on the fins in cryptically-patterned species is hypothesized to provide a visual aid and may function in intraspecific signaling [18,19]. Chlopsid and moray eels are also capable of visualizing their bright green fluorescent emissions, which are produced by fluorescent proteins derived from fatty acid binding proteins [20–22]. In addition, many reef lineages possess yellow intraocular filters in their lenses or corneas [23]. These filters can function as longpass filters, which may enhance the perception of longer wavelength fluorescent emissions within a primarily blue ambient environment [1]. Although further studies are needed to determine and compare the diversity of visual spectral ranges broadly among teleosts, in general, many reef fishes are capable of visualizing the green and/or red portions of the spectrum common in fluorescent emissions [13,16,17].

Several recent studies have documented the widespread presence of biofluorescence across ray-finned fishes [1,8,10,11,19], but relatively few of these report fluorescent emission spectra [1,8,18,19,24]. Anthes et al. [18] documented variation in red fluorescent emissions within several teleost families, finding three general groups of emission peaks: near red, deep red, and far red. However, their analysis excluded all emission peaks below 580 nm (i.e., green-yellow) [18]. No study to date has focused on investigating variation in fluorescent emission peaks across the full range of fluorescent wavelengths observed in teleosts [1,24]. This highlights a large gap in knowledge, especially considering prior studies have reported the presence of multiple fluorescent colors (e.g., green and red) even within an individual [1,9,24].

In this study, we compare emission spectra within eighteen biofluorescent families across the phylogeny of Teleostei [1,9]. We document large variation among families and genera, and over body anatomy within individual species. The objectives of this study are to: 1) record detailed fluorescence emission spectra across a wide array of teleost families that have been shown to exhibit biofluorescence; 2) determine and characterize how fluorescence emission spectra vary within a family, genus, and species; 3) analyze variation in fluorescence emission wavelengths by anatomical region within an individual; and 4) discuss the implications of variation in fluorescent emission spectra (i.e., the presence of several distinct emission peaks within a lineage or species) as it relates to the potential for a diverse array of fluorescent emission patterns or signals to be produced. Our results show that fluorescent emission spectra are diverse both among teleost families, as well as within genera. We also show that fluorescent emissions often vary largely among specific body regions within an individual. This study expands our understanding of the diversity of biofluorescence in teleosts and provides the groundwork for future studies focused on investigating the function of biofluorescence in marine fishes.

## Materials and Methods

### Specimens and imaging

Specimens used in this study are housed at the American Museum of Natural History (AMNH), New York, and are from collections made in the Solomons Islands (2012, 2013, and 2019), Greenland (2019), and Thailand (2024) (S1 Table). Research, collecting, and export permits were obtained from the Ministry of Fisheries and Ministry of Environment (Honiara, Solomon Islands), local fisheries authorities in Greenland, and the Department of Fisheries and Chulalonghorn University (Bangkok, Thailand). This study was carried out in strict accordance with the recommendations in the Guidelines for the Use of Fishes in Research of the American Fisheries Society and the American Museum of Natural History’s Institutional Animal Care and Use Committee (IACUC).

For fluorescence imaging, live and frozen specimens were placed in a narrow photographic tank and gently held flat against a thin glass front. Fluorescent emissions were imaged while in a dark room using a Nikon D800 or D4 DSLR camera outfitted with a Nikon 60 or 105 mm macro lens, or a Sony A7SII or A7RV camera outfitted with a Sony 90 mm macro lens. Flashes (Nikon SB910) were covered with blue interference bandpass excitation filters (Omega Optical, Inc., Brattleboro, VT; Semrock, Inc., Rochester, NY) to elicit fluorescence and longpass (LP) emission filters (Semrock, Inc.) were attached to the front of the camera lens to block any blue excitation light and record only emitted fluorescence. To best capture the fluorescent emissions, multiple LP filter pairs were tested. For example, a 514 nm LP filter was used to capture green fluorescence, whereas a 561 nm LP filter was used to image longer-wavelength fluorescence (orange and red) and to block any emitted green fluorescence in species exhibiting multiple fluorescent colors (i.e., overlapping fluorescent wavelengths).

### Fluorescent spectra measurements

Emission spectra were recorded while in a dark room by using an Ocean Optics USB2000+ portable spectrophotometer (Dunedin, FL) equipped with a hand-held fiber optic probe (Ocean Optics ZFQ-12135). To provide excitation light to elicit fluorescence, specimens were illuminated with Royal Blue LED lights (Pierce Lab, Yale University, New Haven, CT), collimated to ensure perpendicular incidence on the scientific grade 450-470 nm interference filter surface, thereby minimizing the transmission of out-of-band energy. Emission spectra were then recorded by placing the fiber optic probe proximate to specific anatomical parts of the individual fish specimen exhibiting biofluorescence. This process was repeated several times for each specimen and each anatomical region to ensure the accuracy and repeatability of the spectrophotometer readings. To visualize and record longer (i.e., yellow-orange) wavelength emissions in members of Chlopsidae, which exhibit bright green fluorescence over their entire body, it was necessary to use a LP emission filter (561 nm) to block these green fluorescent emissions (see [1]; Fig S4).

Fluorescent emission peaks (lambda-max) are defined as the wavelengths that correspond to the highest intensity value (S2 Fig). When a single spectrum reading exhibited multiple distinct emission peaks (e.g., one green peak and another red peak), peak values were reported for each distinct emission wavelength. All resulting emission spectra were graphed in R [25] using the package “ggplot2” [26] and lines were smoothed in Adobe Illustrator to decrease unnecessary noise without altering data. For some spectra plots, only the emission readings with the highest intensity value were included when redundant lower intensity peaks were present. Plots for all spectra readings taken can be found in supplemental materials (Figs S4-22). The scatter plot of peak values for each family (Fig 1A) was plotted in R [25] using the package “ggplot2” [26]. Visual pigment data (S3 Table) were collected from the literature [27,28] and plotted in R [25] using the package “ggplot2” [26]. Families are listed by taxonomic convention throughout, such that families within specific orders appear sequentially.

**Fig 1.**
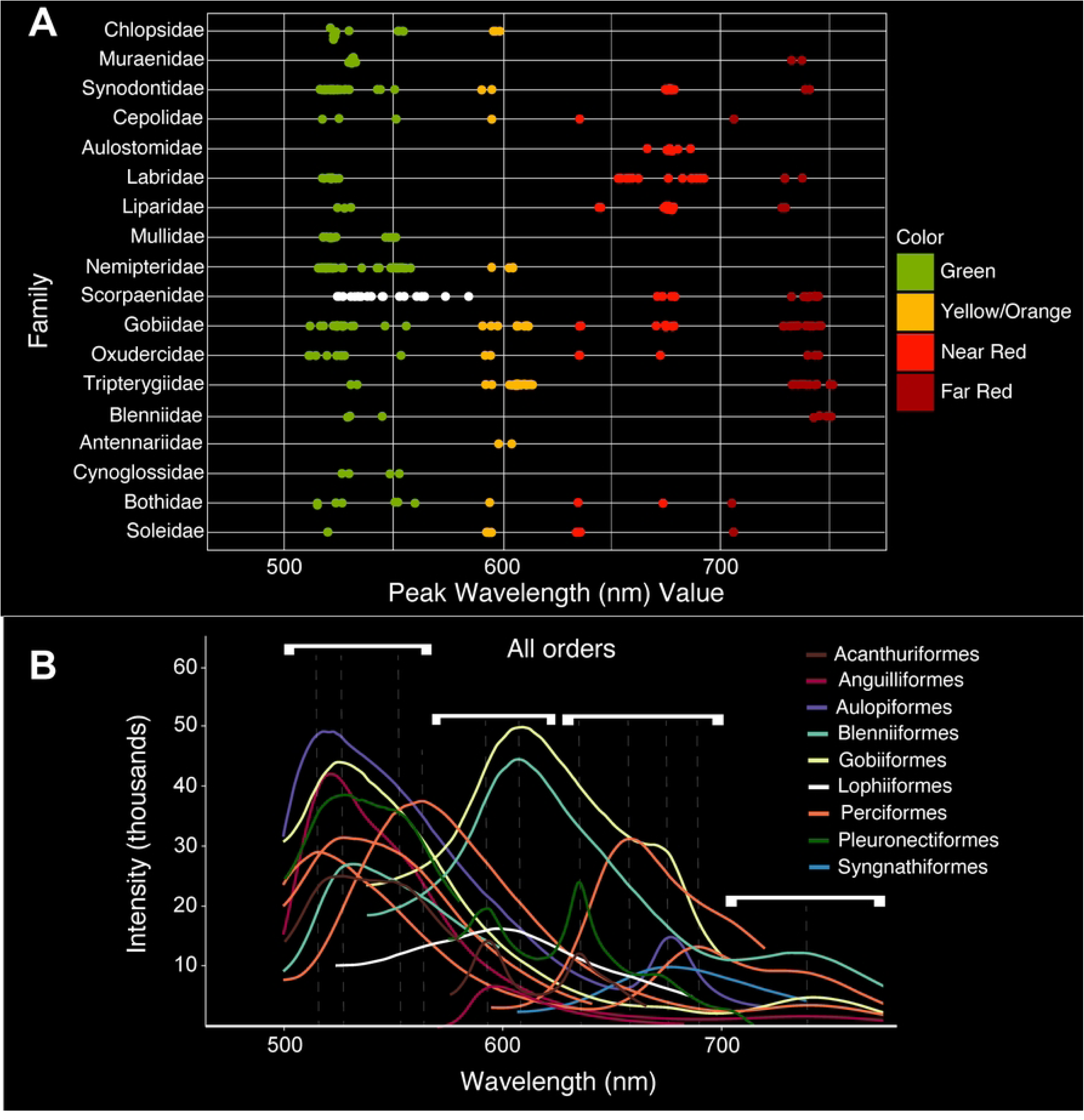
Fluorescent emission spectra of all lineages investigated. A) Emission peaks recorded from species representing 18 teleost families. B) Corresponding spectra recorded for all nine teleost orders investigated: Acanthuriformes (Cepolidae, Nemipteridae), Anguilliformes (Chlopsidae, Muraenidae), Aulopiformes (Synodontidae), Blenniiformes (Blenniidae, Tripterygiidae), Carangiformes (Bothidae, Cynoglossidae, Soleidae), Gobiiformes (Gobiidae, Oxudercidae), Lophiiformes (Antennariidae), Perciformes (Labridae, Liparidae, Scorpaenidae), and Syngnathidae (Aulostomidae, Mullidae). Dashed lines represent fluorescent emission peaks. Intensity values are relative and do not signify overall brightness. White brackets correspond to the four main wavelength/color ranges.

## Results

### Variation in fluorescence among families

In general, four distinct, non-overlapping wavelength (i.e., color) ranges of fluorescent emission peaks are observed among the teleost families we investigated. These peaks correspond to the green (∼500-565 nm), yellow-orange (∼565-625 nm), near red (∼625-700 nm), and far red (∼700-750 nm) portions of the visible spectrum (Figs 1, 2; Table 1). Fluorescent emission peaks ranging from 514-560 nm (green) are present in Chlopsidae, Muraenidae, Synodontidae, Cepolidae, Labridae, Liparidae, Mullidae, Nemipteridae, Scorpaenidae, Gobiidae, Oxudercidae, Tripterygiidae, Blenniidae, Cynoglossidae, Bothidae, and Soleidae. Emission peaks ranging from 573-613 nm (yellow-orange) are present in Chlopsidae, Synodontidae, Cepolidae, Nemipteridae, Scorpaenidae, Gobiidae, Oxudercidae, Tripterygiidae, Antennariidae, Bothidae, and Soleidae. Near red emission peaks ranging from 633-692 nm are observed in Synodontidae, Cepolidae, Aulostomidae, Labridae, Liparidae, Scorpaenidae, Gobiidae, Oxudercidae, Bothidae, and Soleidae. We also find additional far-red emission peaks ranging from 705-751 nm in Muraenidae, Synodontidae, Cepolidae, Labridae, Liparidae, Scorpaenidae, Gobiidae, Oxudercidae, Tripterygiidae, Blenniidae, Bothidae, and Soleidae.

**Fig 2.**
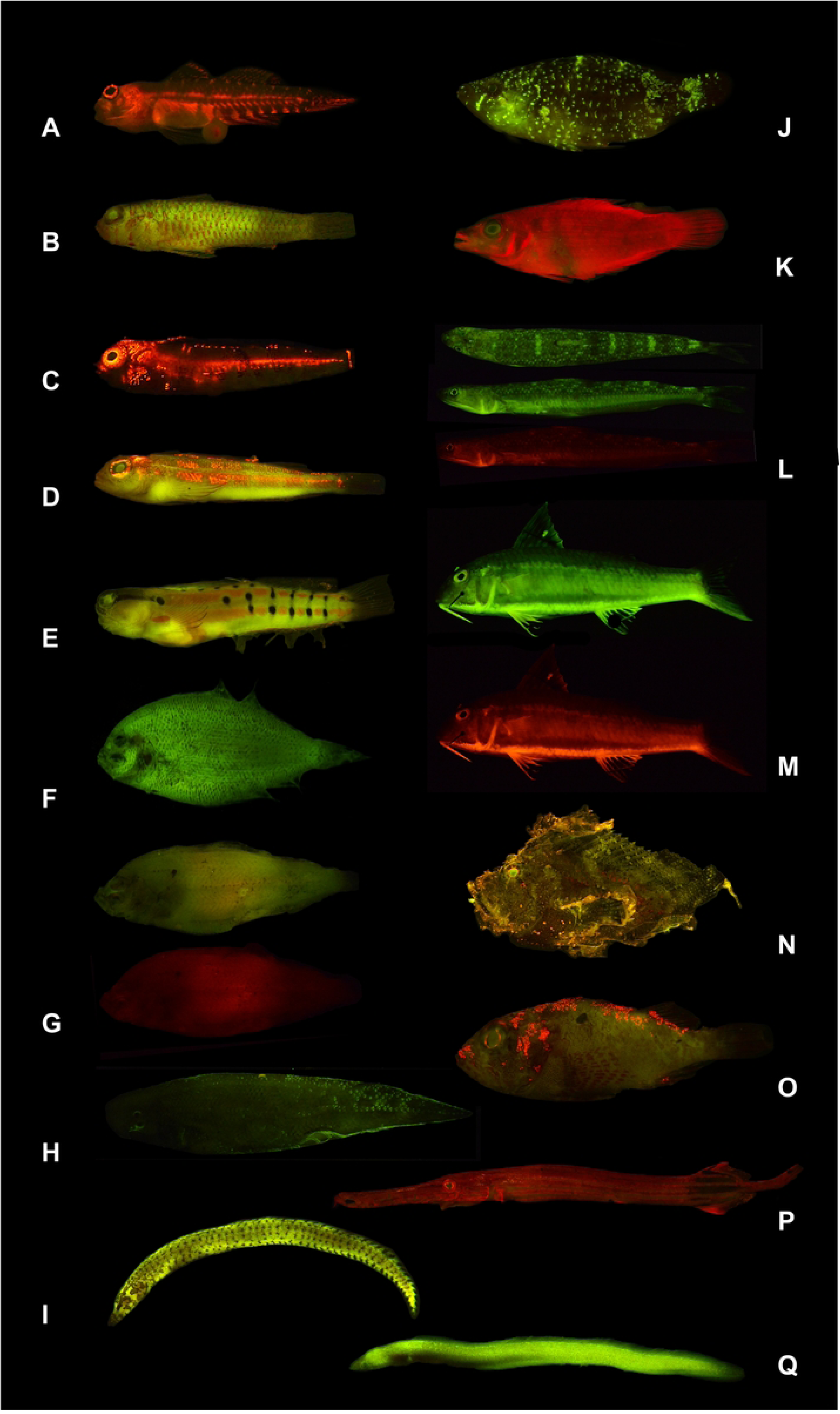
Representative images of biofluorescent teleosts examined in this study. A) *Eviota prasites* (Gobiidae), B) *Trimma fangi* (Gobiidae), C) *Enneapterygius niger* (Tripterygiidae), D) *Helcogramma striata* (Tripterygiidae), E) *Ecsenius axelrodi* (Blenniidae), F) *Engyprosopon mozambiqense* (Bothidae), G) *Japonolaeops dentatus\** (Bothidae), H) *Cynoglossus microlepis* (Cynoglossidae), I) *Gymnothorax zonipectis* (Muraenidae), J) *Cheilinus oxycephalus* (Labridae), K) *Pseudocheilinus evanidus* (Labridae), L) *Saurida micropectoralis*\* (Synodontidae), M) *Upeneus sundaicus\** (Mullidae), N) *Taenianotus triacanthus* (Scorpaenidae), O) *Sebastapistes fowleri* (Scorpaenidae), P) *Aulostomus chinensis* (Aulostomidae), Q) *Kaupichthys diodontus* (Chlopsidae). *Images of the same specimen without (top) and with (bottom) a 561 nm longpass filter to block green fluorescence. *Saurida micropectoralis* (L) is shown in dorsal and lateral views for green fluorescence.

**Table 1.** General ranges of fluorescent emission spectra peaks (nm) in all teleost families investigated.

| Family | Green (~500-565 nm) | Yellow-Orange (~565-625 nm) | Near Red (~625-700 nm) | Far Red (~700-750 nm) | Eye Range (nm) |
| --- | --- | --- | --- | --- | --- |
| Chlopsidae | 520-528, 551-554 | 596-599 |  |  | 520-522, 554 |
| Muraenidae | 529-532 |  |  | 732-736 |  |
| Nemipteridae | 516-558 | 594-604 |  |  | 518-521, 552-555 |
| Cepolidae | 516-525, 551 | 594 | 635 | 706 | 525, 551 |
| Synodontidae | 516-529, 542-550 | 590-594 | 674-678 | 738-740 | 518-529, 542-543 |
| Aulostomidae |  |  | 666-686 |  | 680 |
| Mullidae | 517-523, 547-550 |  |  |  | 520-523, 546-550 |
| Blenniidae | 529-545 |  |  | 743-751 |  |
| Tripterygiidae | 530-532 | 592-613 |  | 732-751 | 605-608 |
| Labridae | 517-524 |  | 652-692 | 729-737 |  |
| Scorpaenidae | 524-554 | 573 | 673-678 | 732-740 | 524-530, 677, 732 |
| *Sebastapistes* | 526-584 |  | 670 | 737-744 | 526-544, 584, 742 |
| Liparidae | 524-530 |  | 643-645, 673-677 | 726-729 | 674-677 |
| Gobiidae | 514-531, 546-555 | 590-611 | 635, 652-678 | 731-745 | 525-527, 546-551 |
| Oxudercidae | 514-527, 553 | 592-594 | 635-672 | 706, 739-744 |  |
| Antennariidae |  | 598-604 |  |  |  |
| Bothidae | 515-526, 551-560 | 594 | 635, 674 | 705 | 560 |
| Soleidae | 519 | 592-594 | 633-635 | 705 |  |
| Cynoglossidae | 526-529, 548-552 |  |  |  |  |

Emission peaks are defined as the wavelength associated with the greatest intensity value for each distinct emission curve.

Members of Synodontidae, Cepolidae, Scorpaenidae, Gobiidae, Oxudercidae, Bothidae, and Soleidae exhibit fluorescent emissions in all four of these wavelength ranges (Fig 1; Table 1). However, the scorpaenid genus *Sebastapistes* is an exception to the distinct wavelength ranges identified above and exhibits a continuous range of fluorescent emission peaks from 524-583 nm, spanning wavelengths that correspond to green and yellow-orange (Figs 1 and 3C; Table 1). Nine of the 18 families examined were found to exhibit at least four distinct fluorescent emission peaks, including members of Synodontidae, Cepolidae, Labridae, Liparidae, Scorpaenidae, Gobiidae, Oxudercidae, Bothidae, and Soleidae (Figs 1 and 3; S2, S8, S11, and S21 Figs).

**Fig 3.**
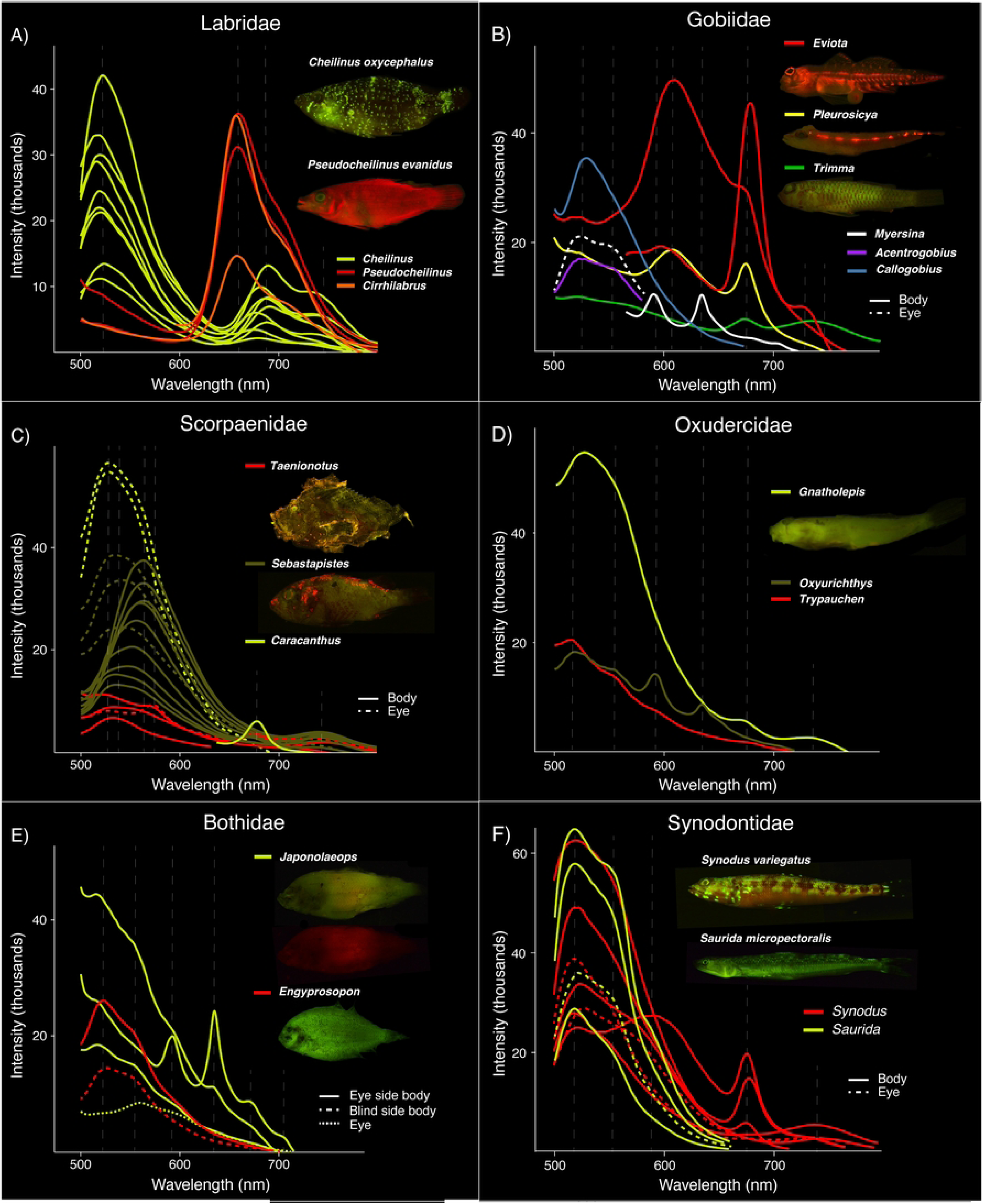
Fluorescent emission spectra recorded from various regions of the body. Variation among genera is shown for members of: A) Labridae; B) Gobiidae; C) Scorpaenidae; D) Oxudercidae; E) Bothidae; and F) Synodontidae. Dashed lines represent fluorescent emission peaks.

### Dual fluorescent emission peaks

Members of some biofluorescent families exhibit dual emission peaks, where a spectral reading shows two distinct emission peaks within a small wavelength (i.e., single color) range (e.g., Figs 1B Pleuronectiformes and 3B *Myersina*). Dual green emission peaks are present in members of Chlopsidae (520-528 and 551-554 nm; S4 Fig), Cepolidae (525 and 550 nm; S8 Fig), Mullidae (517-523 and 547-550 nm; S12 Fig), Nemipteridae (517-536 and 549-558 nm; S13 Fig), Gobiidae (523-527 and 546-555 nm; Fig 3B), Cynoglossidae (526-529 and 548-552 nm; S20 Fig), and Bothidae (523-526 and 551-552 nm; S21 Fig). Dual emission peaks within the near red portion of the spectrum are only present in Liparidae (643-645 and 676 nm; S11 Fig) and Bothidae (635 and 674 nm; S21 Fig).

### Variation within families

Large variation is present in fluorescent emission peaks among genera within Synodontidae, Labridae, Scorpaenidae, Gobiidae, Oxudercidae, and Bothidae. In Synodontidae, green fluorescent emissions are present as a single peak in *Synodus* (516-529 nm) and as a dual peak in *Saurida* (517-525 and 542-550 nm) (Figs 1 and 3F; Table 1). Yellow-orange fluorescent emissions produce a prominent peak in *Synodus* (594 nm) and a much weaker peak in *Saurida* (590 nm). Only *Synodus* has fluorescent emissions corresponding to near red (674-678 nm) and far red (738-670 nm) wavelengths; red fluorescent emissions are absent in *Saurida* (Fig 3F). Within Labridae, fluorescent emission peaks in *Cheilinus* are recorded at 517-524 nm (green), 675-692 nm (near red), and 729-737 nm (far red), whereas *Cirrhilabrus* and *Pseudocheilinus* both exhibit near red peaks at 652-662 nm (Figs 1 and 3A; Table 1) and lack green fluorescent emissions.

In Scorpaenidae, all genera examined exhibit green fluorescent emission peaks (Figs 1 and 3C; Table 1). These peaks range from 526-530 nm in *Caracanthus* and 524 nm in *Taenianotus*. *Sebastapistes* exhibits a wide range of emission peaks spanning wavelengths from green to yellow-orange (526-584 nm). However, a distinct yellow emission peak is present in *Taenianotus* at 573 nm. Near red emission peaks are present in *Caracanthus* (679 nm), *Sebastapistes* (670 nm), and *Taenianotus* (673-677 nm), whereas far red emission peaks are present in *Sebastapistes* (717-744 nm) and *Taenianotus* (732-740 nm) (Fig 3C).

Considerably more emission peak variation is recorded among genera in Gobiidae (Figs 1 and 3B; Table 1). Green fluorescence is present in *Callogobius* (526-531 nm), *Eviota* (524 nm), and *Trimma* (517-526 nm) as a single peak, and as a dual peak in *Acentrogobius* (523 and 555 nm) and *Myersina* (517-527 and 546-555 nm). Yellow-orange fluorescent emissions are present as a single peak in *Eviota* (594-611 nm), *Myersina* (590 nm), and *Pleurosicya* (606-611 nm). Near red emission peaks are present in *Eviota* (672-678 nm), *Myersina* (635 nm), *Pleurosicya* (674 nm), and *Trimma* (670-674 nm). Far red fluorescent emissions are present in *Eviota* (728-731 nm) and *Trimma* (731-745 nm). In the gobiiform family Oxudercidae, green fluorescence is present as a single peak in both *Gnatholepis* (524-527 nm) and *Trypauchen* (517 nm), and as a dual peak in *Oxyurichthys* (514-519 and 553 nm) (Figs 1 and 3D; Table 1). Yellow-orange fluorescence is present as a single peak in *Oxyurichthys* (592-594 nm). Near red fluorescent emissions are present in both *Gnatholepis* (672 nm) and *Oxyurichthys* (635 nm), however, far red fluorescence is only present in *Gnatholepis* at (739-744 nm) (Fig 3D).

In Bothidae, green fluorescent emissions are present as a single peak in *Japonolaeops* (515 nm) and as a dual peak in *Engyprosopon* (523-526 and 551-552 nm) (Figs 1 and 3E; Table 1). Yellow-orange, near red, and far red fluorescent emissions are only present in *Japonolaeops* at 594 nm, 635 and 674 nm (dual emission), and 705 nm, respectively (Fig 3E).

We find little to no variation in fluorescent emission wavelength peaks among the two genera of Tripterygiidae (*Enneapterygius* and *Helcogramma*), Nemipteridae (*Nemipterus* and *Scolopsis*), or Soleidae (*Heteromycteris* and *Zebrias*) that we examined in this study (S17 and S13 Figs). In the remaining families investigated, we were only able to analyze a single genus (Chlopsidae, Muraenidae, Cepolidae, Aulostomidae, Liparidae, Mullidae, Blenniidae, Antennariidae, and Cynoglossidae) so intrafamilial variation could not be assessed (S4, S5, S8, S9, S11, S12, and S18-20 Figs).

### Eye fluorescent emissions

Fluorescence from the eye (iris and lens), is commonly observed in marine teleosts (Fig 2). Fluorescent emission spectra were recorded from the eyes in members of eleven teleost families (Fig 4; Table 1). We recorded a similar double green emission peak from the eyes of members of Chlopsidae (520-522 and 554 nm), Synodontidae (518-529 and 542-543 nm), Cepolidae (525 and 551 nm), Mullidae (520-523 and 546-550 nm), Nemipteridae (518-521 and 552-555 nm), and Gobiidae (*Myersina*, 525-527 and 546-551 nm). Green emissions from the eyes are also present as a single peak in Scorpaenidae (524-544 nm) and Bothidae (*Japonolaeops,* 560 nm). In the scorpaenid genus *Sebastapistes,* we recorded a yellow-orange emission peak from the eyes at 584 nm, whereas in the tripterygiid genus *Helcogramma,* a yellow-orange emission peak was recorded at 605-608 nm (Fig 4). Aulostomidae, Liparidae, and Scorpaenidae (*Taenionotus*) have eyes that emit near red peaks at 680 nm, 675 nm, and 677 nm, respectively. The eyes of members of Scorpaenidae also emit a far red emission peak at 732-742 nm.

**Fig 4.**
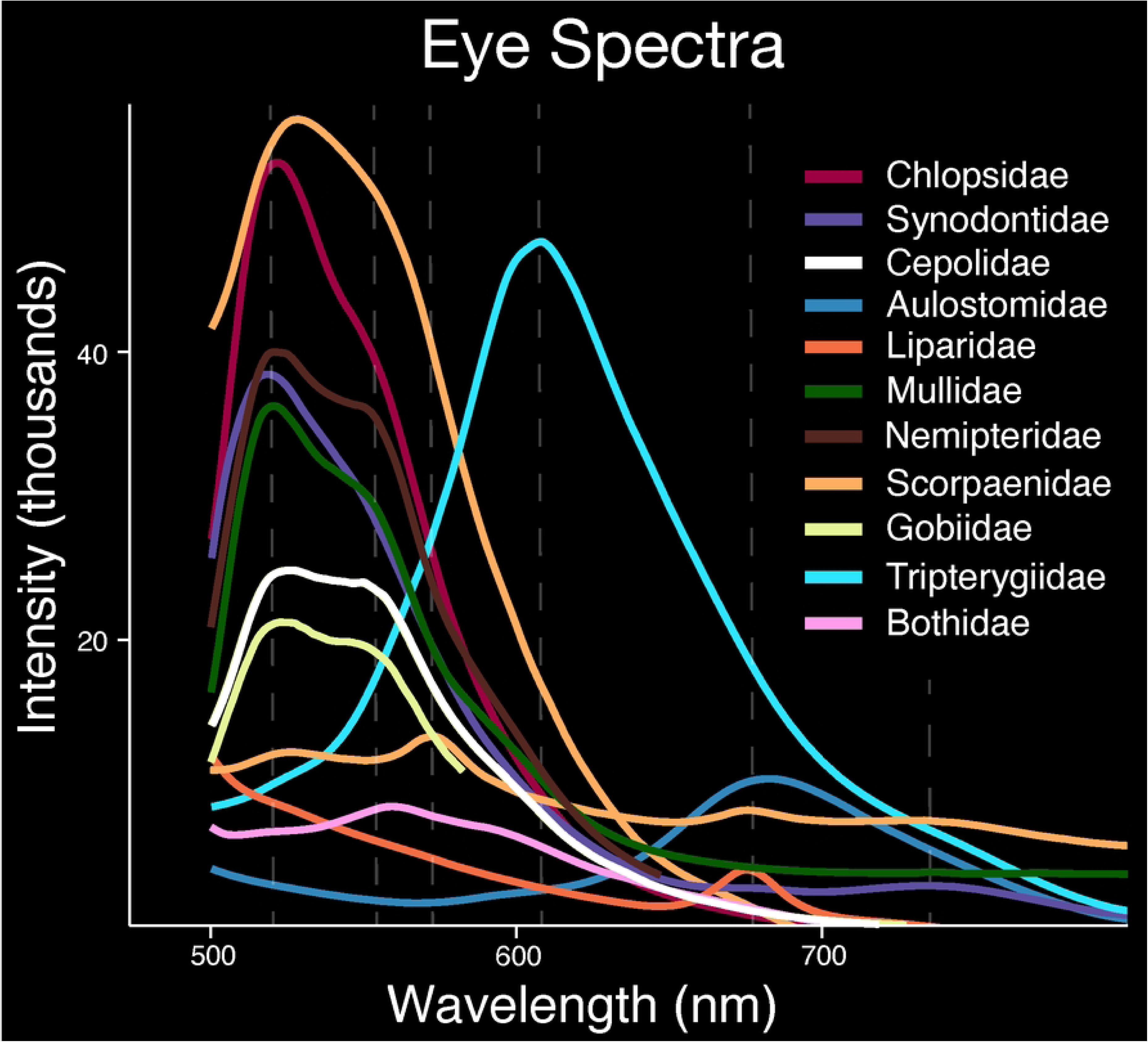
Fluorescent emission spectra recorded from the eyes. Eleven teleost families were measured: Chlopsidae, Synodontidae, Cepolidae, Aulostomidae, Liparidae, Mullidae, Nemipteridae, Scorpaenidae, Gobiidae, Tripterygiidae, and Bothidae. Dashed lines represent fluorescent emission peaks.

### Intraspecific fluorescent emission variation by anatomical region

In some individuals, fluorescent emissions vary by body region (Fig 5; Table 1). In *Helcogramma striata* (Tripterygiidae), the upper and lower flank exhibit a green emission peak ranging from 530-532 nm, whereas only the eye and portion of the upper flank share a similar yellow-orange emission peak of 594-608 nm (Fig 5A). The lower flank of *H. striata* also has far red emission peaks ranging from 733-751 nm. In Scorpaenidae, *Taenionotus triacanthus* exhibits four distinct emission peaks over different body regions, including the eye, mouth, and upper flank (Fig 5B; Table 1). Emission peaks are green (524 nm) for the mouth and eye, yellow-orange (573 nm) for the mouth, eye, and entire flank, near red (673-677 nm) for the mouth and eye, and far red (732-740 nm) for the upper flank and eye. In *Sebastapistes strongia* (Scorpaenidae), the eye exhibits a green emission peak at 544 nm and a single broad yellow-orange peak at 584 nm, spanning ∼566-596 nm (Fig 5C). The upper flank exhibits a green emission peak at 560-564 nm and an additional far red peak in the same spectra reading at 737-743 nm (Fig 5C). The anterior and posterior body of one *Gymnothorax zonipectis* (Muraenidae) specimen exhibits a similar green emission peak from 529-532 nm, however, the posterior body has an additional far red peak from the same spectra reading at 732-736 nm (Fig 5D). In *Cepola schlegelii* (Cepolidae), green emission peaks are present in the eyes (double peak, 525 and 551 nm) and posterior region of the body (516 nm) but are absent in the anterior section of the body (S8 Fig); both the anterior and posterior regions of the body have a yellow-orange emission peak at 594 nm and a near red emission peak at 635 nm. However, only the anterior region of the body of *C. schlegelii* exhibits a far red emission peak at 706 nm (S8 Fig). All other specimens examined show little to no variation over the various body regions that were scanned for fluorescence.

**Fig 5.**
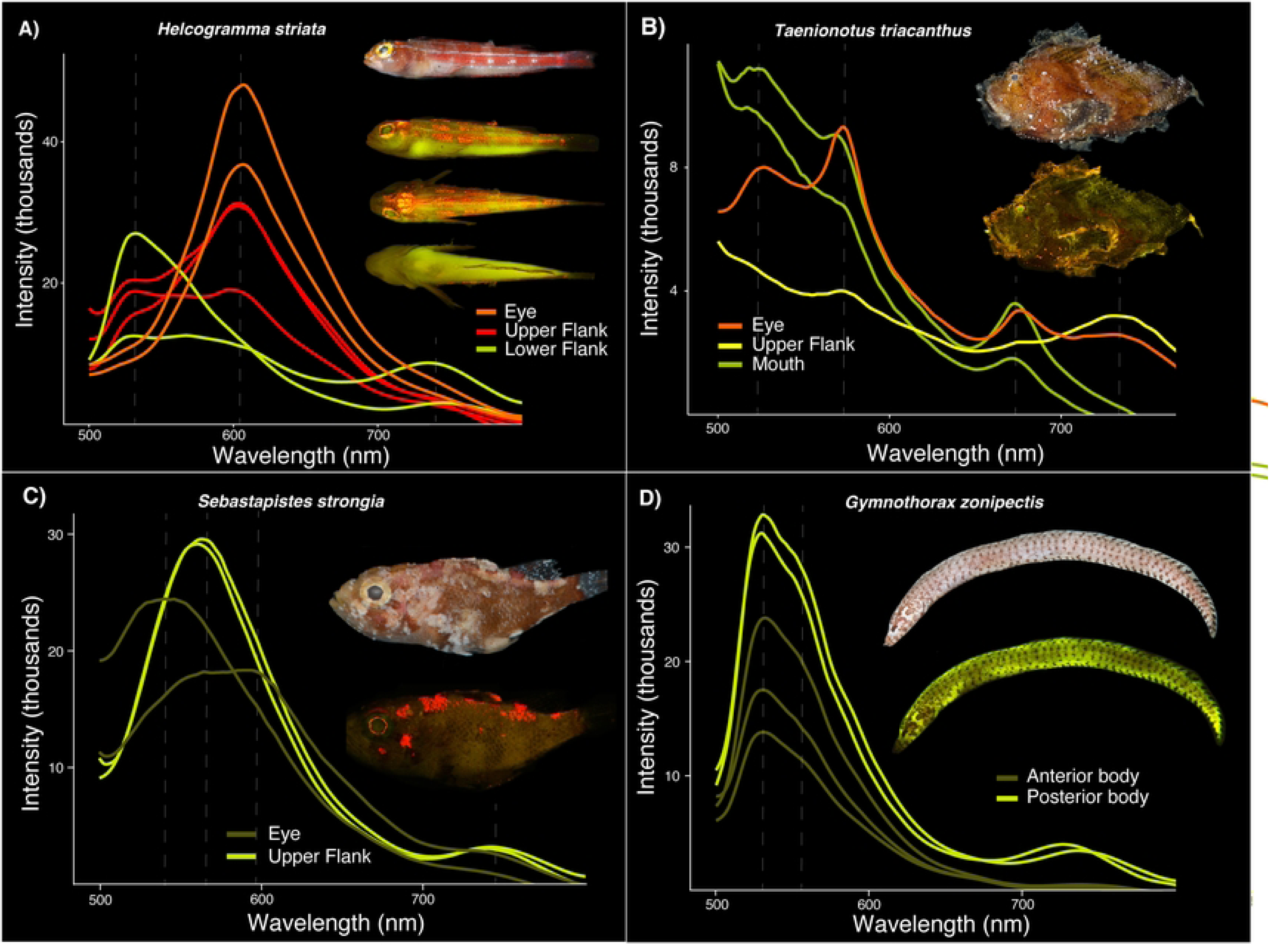
Variation in fluorescent emission spectra, or lack thereof, over different regions of the body in individuals. A) *Helcogramma striata* (Tripterygiidae); B) *Taenianotus triacanthus* (Scorpaenidae); C) *Sebastapistes strongia* (Scorpaenidae); D) *Gymnothorax zonipectis* (Muraenidae). Note: For each panel the top image shows the species imaged under white light and all others show the species fluorescing. Dashed lines represent fluorescent emission peaks.

### Visual pigments

Visual pigment data is available for species representing nine of the 18 families examined in this study (Fig 6; S3 Table) [27,28]. All families have visual pigments with peak absorbances (lambda max) within the 366-500 nm range. Visual pigments with peak absorbances in green wavelengths are present in Muraenidae (510 nm), Synodontidae (503 nm), Labridae (505-555 nm), Liparidae (522-527 nm), Mullidae (515-533 nm), Scorpaenidae (501-530 nm), Gobiidae (508-553 nm), and Blenniidae (500-561 nm). Visual pigments with peak absorbances in yellow-orange wavelengths are present in Gobiidae (565 nm) and Blenniidae (570 nm).

**Fig 6.**
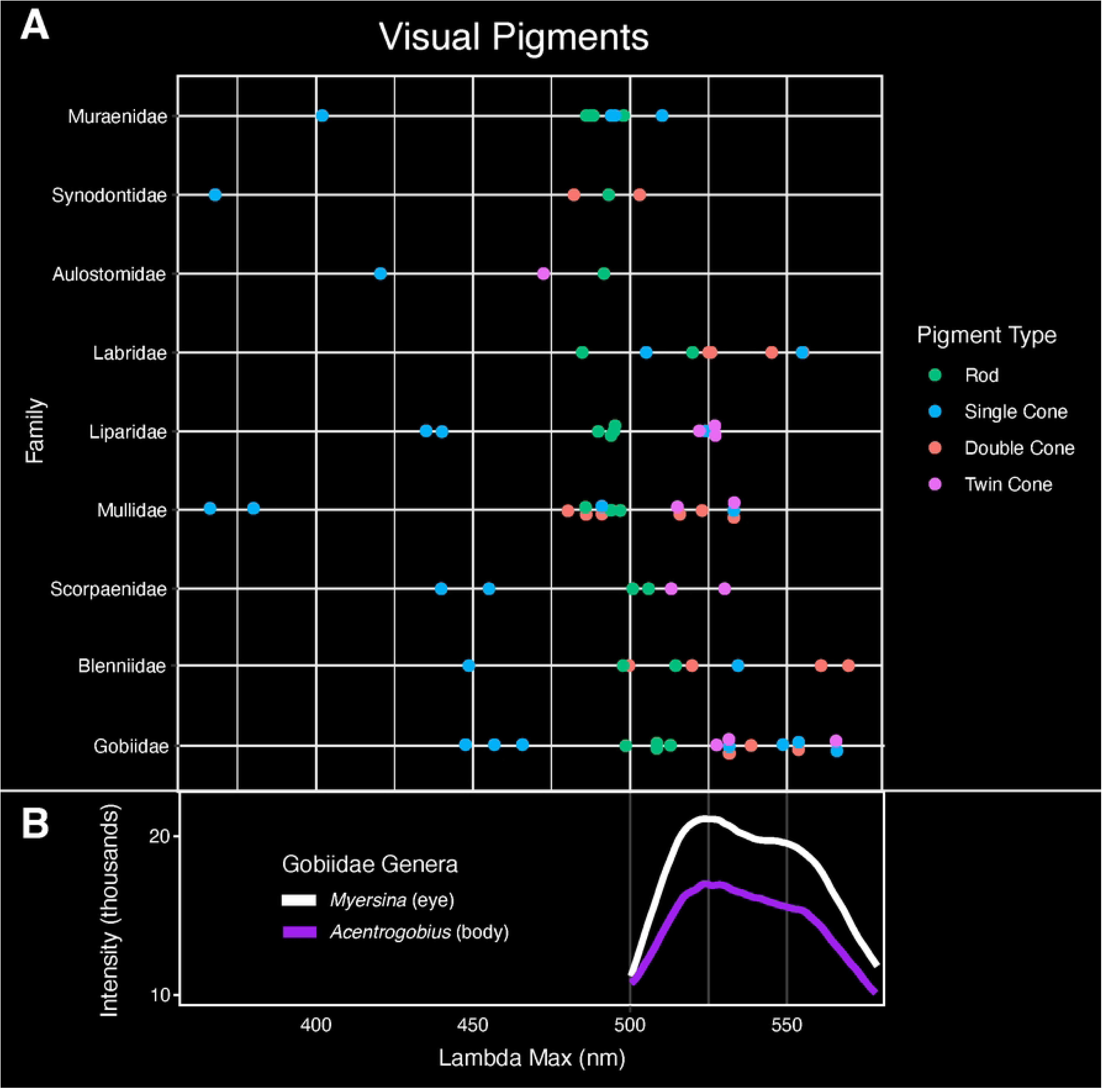
Visual pigments of nine families investigated in this study. Peak absorbance (lambda max) is reported for each pigment type: rod, single cone, double cone, and twin cone.

## Discussion

In this study we compare the emission spectra of numerous species representing 18 diverse teleost families that exhibit biofluorescence. Recent studies have shown that biofluorescence is phylogenetically widespread and phenotypically variable in marine fishes, particularly across ray-finned lineages, where fluorescent emissions have generally been reported simply as either red or green [1,9]. Our results, however, show that fluorescence is far more variable in emitted wavelength and corresponding color, both among and within lineages at the family, genus, and species scale (Figs 1-5; Table 1). Further, in many species, fluorescent emissions vary within an individual (Fig 5; Table 1). In general, we recover a pattern of four distinct emission peak ranges spanning green (514-560 nm), yellow-orange (590-613 nm), near red (633-692 nm), and far red (705-751 nm) wavelengths (Fig 1; Table 1). Interestingly, fluorescent emissions in yellow wavelengths (∼565-590 nm) are quite rare, only occurring in two scorpaenid genera, *Sebastapistes* (584 nm) and *Taenianotus* (573 nm) (Fig 3C; Table 1; S2 Table).

Overall, we find that fluorescent emissions are highly variable in marine teleosts. For example, three families-Gobiidae, Oxudercidae, and Bothidae - exhibit at least six distinct, non-overlapping fluorescent emission peaks (Figs 1 and 3; Table 1). A lack of variation in fluorescent emissions was rare, and only two of the 18 families examined exhibited a single fluorescent peak: yellow-orange in Antennariidae and near red in Aulostomidae (Fig 1; Table 1). All other families examined had at least two distinct fluorescent emission peaks, with some falling within the same emission wavelength range (e.g., green) (Fig 1; Table 1). We note that previous studies have reported fluorescence in other wavelength ranges (colors) in members of both Antennariidae, and Mullidae [1,9,18], in addition to the spectra reported herein.

### Vision

In the marine environment, fluorescence may function by either enhancing the color signal (i.e., specific emission wavelengths are visible) or increasing luminosity contrast with the surrounding background [11,19]. Longer wavelengths are rapidly absorbed in oceanic habitats and fluorescence can restore these lower-energy, longer wavelengths (green-red) at depths where only shorter blue wavelengths can penetrate [1,11]. Reef fishes generally have good color vision in green and/or red wavelengths, including many of the families examined this study (e.g., Gobiidae, Labridae) [13,16,17]. Species representing nine of the families investigated in this study had available visual pigment data in the literature (Fig 6; S3 Table) [27,28]. Except for Aulostomidae, all families had visual pigments with peak absorbances (lambda max) at similar green wavelengths to their fluorescent emissions (500-565 nm; Figs 1 and 6A). Gobiidae in particular had visual pigments with peak absorbances that closely matched the green fluorescent dual emission peaks found in *Myersina* and *Acentrogobius* (Fig 6B). However, three families we examined, Blenniidae, Gobiidae, and Labridae, had particularly longer wavelength shifted vision, with peak absorbances extending to 570 nm (Fig 6A). Whereas peak absorbances are the optimal wavelength for photon capture of each visual pigment, the spectral sensitivity curves for many marine species extend well beyond these optimal wavelengths [8,13,17].

In addition, members of several reef families (e.g., Pomacentridae, Gobiidae, Labridae) possess long wavelength sensitivity (LWS) opsins in their eyes, allowing them to visualize red wavelengths [13,16,17]. For example, *Eviota pellucida* (Gobiidae), a species that emits red fluorescence, has a spectral range that extends to ∼650 nm but with a spectral sensitivity that peaks at 450-550 nm [8]. Moreover, many lineages of reef associated fishes (e.g., Synodontidae, Labridae, Scorpaenidae, Pleuronectiformes) that exhibit fluorescence have been shown to possess yellow intraocular (lenses or cornea) filters [23] that absorb short wavelengths of light. These filters could enable enhanced perception of longer wavelength fluorescent emissions in a blue ambient environment.

While some fluorescent wavelengths (i.e., far red) may not be associated with a color signal in some fishes, longer wavelengths may function to create greater luminosity contrast with surrounding backgrounds in a monochromatic blue ambient environment at depth, as shown in catsharks [11]. In dimmer oceanic waters (e.g., turbid coastal habitats, deeper environments where only blue light penetrates), emitted fluorescent photons at longer wavelengths may function to enhance luminosity contrast in the individual [11]. The particular functionality of this enhancement depends largely on the spectral sensitivity and environment of the signal receiver (see Marshall and Johnsen[29] for specific criteria). For example, the swell shark (*Cephaloscyllium ventriosum*) has only a single visual pigment and lacks pre-retinal filters. As a result, detecting a fluorescent pattern against a background must rely entirely on variation in luminosity (i.e., brightness). Although the visual pigment (maximum absorbance 484 nm +/-3 nm) in the swell shark is not spectrally situated to maximize contrast between the individual’s bright green fluorescence and the blue background light, the absorbance of their visual pigment overlaps the spectral bandwidth of their green fluorescence. As a result, the reticulated patterns of dark and light on the body can still be detected [11]. Luminosity contrast resulting from fluorescence in *C. ventriosum* can result in a conspecific being more apparent than a non-fluorescent shark [11]. Thus, fluorescence can serve a functional role in luminosity contrast, even if fluorescent wavelengths (i.e., far red) are beyond the color signal range of an organism. Consequently, far red fluorescence should not be discounted as nonfunctional without further testing of its potential role in luminosity contrast.

In addition to serving as an added visual signal for conspecifics, as has been shown for gobies (Gobiidae) and triplefins (Tripterygiidae) [11], fluorescence in the eyes of some reef fishes may also enhance prey detection[18,19]. In *Tripterygion delaisi* (Tripterygiidae), fluorescence in the iris (∼600 nm) may induce high contrast reflections in the eyes of cryptic prey, increasing conspicuousness and thus capture success [30]. Similar mechanisms are hypothesized to be used by deep-sea dragonfishes (Stomiiformes) and flashlight fishes (Anomalopidae) [31,32]. Whereas we find comparable orange wavelengths of fluorescence in the eyes of members of Tripterygiidae (605-608 nm), we also find longer wavelengths of near red fluorescence in the eyes of members of Aulostomidae, Liparidae, and Scorpaenidae (near red, 674-680 nm) (Fig 4; Table 1). These red fluorescent emissions are more similar in wavelength to those observed in dragonfishes (678 nm), and could possibly serve a similar visual function in prey capture [32]. Thus, fluorescence can potentially serve multiple visual functions in both conspecific identification through contrast enhancement and prey detection [11,18,19].

### Variation in fluorescence emissions

It is currently unknown whether slight variations in fluorescent emission wavelengths could serve a functional role, such as aiding in either intra- or interspecific recognition, camouflage, or by providing unique visual cues. In closely related species where patterns under white light are almost identical, variations in fluorescent emissions may provide an extra visual cue to aid in species recognition (e.g., reef lizardfishes, Synodontidae; see [1] Fig 3). In Synodontidae, *Synodus* species exhibit both near red and far red emission peaks that are absent in members of the closely related genus *Saurida*. These differences in fluorescent emissions may aid in distinguishing these otherwise morphologically similar genera, supporting the hypothesized function of fluorescence in mate identification/species recognition (Fig 3F). Red fluorescence has also been tied to intraspecific signaling in some cryptic reef fishes [e.g., members of Syngnathidae, Gobiidae (*Eviota*), and Tripterygiidae] when present on certain body parts used in mating (fins) [8]. However, we find no variation in the wavelength of fluorescence emissions over the body in species of Tripterygiidae or *Eviota* that were investigated in this study (S1 and S2 Figs). Thus, the role of fluorescence in intraspecific signaling in these groups could be more dependent on the location of the fluorescent emission rather than emission wavelength itself. However, we did find variation in fluorescent emissions over the body of some individuals (Figs 4 and 5). For example, *Helcogramma striata* (Tripterygiidae) exhibits green and yellow-orange fluorescence over all body regions (eye, upper and lower flank), whereas far red emissions are only present on the lower flank (Fig 5A). Future studies are needed to determine if the behavior and distribution of fluorescent tissue over the body of individuals influences success in intraspecific signaling.

Marine fishes may also use biofluorescence to aid in camouflage. This is likely in some species of Nemipteridae and Scorpaenidae, which have been observed resting or potentially hiding near fluorescent substrates or structures (e.g., algae and corals) with similar fluorescent emissions to themselves [1]. Coral fluorescence mainly falls within green (490-560 nm) or yellow-orange (580-600 nm) wavelengths [33–35], similar to fluorescent emissions present in individuals in both Nemipteridae and Scorpaenidae (Fig 1). Interestingly, the individuals we assessed in the scorpaenid genera *Sebastapistes* uniquely exhibit a continuous range of emission peaks spanning wavelengths from green to yellow-orange (524-583 nm) (Figs 2N-O, 3C, and 5C; Table 1). No species from any other family examined in this study had emission peaks within the 560-590 nm range (Fig 1; Table 1). Algal fluorescence is due to the presence of chlorophyll, which has a near red emission peak at 680 nm, comparable to those seen in seven teleost families measured in this study (652-692 nm) (Fig 1; Table 1) [18]. As a result, fluorescent emissions in teleosts may be multifunctional in intraspecific recognition, camouflage/predator avoidance, and prey detection depending on the location on the body and wavelength of these emissions.

### Fluorescent molecules

To date, molecules and proteins that produce fluorescence in fishes have only been identified and isolated from catsharks and true eels (Anguilliformes), all of which only produce green fluorescence [20–22,36–37]. No molecules or fluorescent proteins that emit yellow through red fluorescent wavelengths have been identified or isolated from fishes, despite the remarkable variation in yellow-orange and near and far red fluorescent emission spectra reported in this study (Fig 1; Table 1). Based on the results from prior studies focused on isolating and characterizing fluorescent proteins from false moray eels, moray eels, and catsharks, fluorescent proteins and other fluorescent molecules in fishes tend to have very narrow emission peaks of around 3 nm maximum [20,22,36–37]. Our results also show that it is quite common for members of several families to emit multiple distinct fluorescent emission peaks within either the green or red (near and far red) portions of the visible spectrum (Fig 1; Table 1). This ability to produce distinct emission signals within a single-color bandwidth greatly increases the variability of potential fluorescent signals that could be produced by an individual.

Our results suggest that either 1) numerous fluorescent proteins or molecules capable of producing fluorescence are present in marine fishes (e.g., multiple green or red fluorescent proteins that result in the dual peaks reported in several families), or 2) that certain lineages can alter emitted wavelengths produced by a single fluorescent protein or molecule. These hypothesized wavelength shifts could be produced via interactions with other pigment producing cells like chromatophores in combination with some type of filter or lens [38], or via some other mechanism leading to a shift in emitted wavelength [39]. Regardless, our results point to a fascinating array of fluorescent colors that can be emitted by marine fishes, frequently within in a single individual, and highlight the need for further investigation of the fluorescent proteins or other molecules responsible for producing these fluorescent emissions.

## Conclusion

Our results show that biofluorescence in teleosts is not only phylogenetically widespread, but phenotypically variable. We find an exceptional degree of variation in fluorescence emission spectra within both families and genera (Figs 1 and 3; Table 1), and even over the body within certain individuals (Figs 4 and 5). This remarkable variation found across a wide array of fluorescent teleost families, could allow for an incredibly diverse and elaborate fluorescent emission signaling system that is highly variable in emitted wavelength, potentially resulting in unique, species-specific fluorescent patterns. Variation in emission spectra could enable marine fishes to produce an exceptionally wide range of visual cues for intraspecific recognition, prey attraction, or predator avoidance/camouflage [1,8,18,19]. Given that marine fishes can produce such a fascinating diversity of fluorescent emission wavelengths (colors), novel studies across a broader range of fluorescent lineages are needed to determine the visual systems/capabilities of additional fluorescent species, identify and isolate the fluorescent molecules capable of producing these stunning displays, and investigate the potential functions of this exceptionally variable phenomenon.

## Acknowledgements

Research, collecting, and export permits were obtained from the Ministry of Fisheries and Ministry of Environment, Honiara, Solomon Islands, from local authorities in Greenland, and from the Department of Fisheries and Chulalonghorn University (CU), Bangkok, Thailand. We are grateful to Somsak Panha and Piyoros (Pomme) Tongkerd (CU) for facilitating our collections and studies in Thailand. We thank Sven Gust of Northern Explorers Diving & Expeditions for SCUBA diving, boat, and logistical support in eastern Greenland. We are grateful to David Gruber (CUNY), Robert Schelly (USFWS), and Brennan Phillips (URI) for considerable assistance in the field and for help with imaging and recording of emission spectra. For assistance with the identification of specimens used in this study we thank Christine Thacker (UCSB), Gerald Allen (WAM), Thomas Monroe (NOAA), and Kunio Amaoka (Hokkaido University).

## Supporting Information Captions

**S1 Table. All specimens used in this study.**

**S2 Table. Fluorescent emission peaks (lambda max) for all specimens.** The unit of peak values are nanometers (wavelength).

**S3 Table. Visual pigments for species representing nine of the 18 families examined in this study.** All data was obtained from the literature [27,28].

**S4 Fig. Fluorescent emission spectra of Chlopsidae.** A longpass emission filter (561 nm) was used to block bright green fluorescent emissions and record yellow-orange wavelengths. Dashed lines represent fluorescent emission peaks.

**S5 Fig. Fluorescent emission spectra of Muraenidae.** Dashed lines represent fluorescent emission peaks.

**S6 Fig. Fluorescent emission spectra of *Saurida* (Synodontidae).** Dashed lines represent fluorescent emission peaks.

**S7 Fig. Fluorescent emission spectra of *Synodus* (Synodontidae).** Dashed lines represent fluorescent emission peaks.

**S8 Fig. Fluorescent emission spectra of Cepolidae.** Dashed lines represent fluorescent emission peaks.

**S9 Fig. Fluorescent emission spectra of Aulostomidae.** Dashed lines represent fluorescent emission peaks.

**S10 Fig. Fluorescent emission spectra of Labridae.** Dashed lines represent fluorescent emission peaks.

**S11 Fig. Fluorescent emission spectra of Liparidae.** Dashed lines represent fluorescent emission peaks.

**S12 Fig. Fluorescent emission spectra of Mullidae.** Dashed lines represent fluorescent emission peaks.

**S13 Fig. Fluorescent emission spectra of Nemipteridae.** Dashed lines represent fluorescent emission peaks.

**S14 Fig. Fluorescent emission spectra of Scorpaenidae.** Dashed lines represent fluorescent emission peaks.

**S15 Fig. Fluorescent emission spectra of Gobiidae.** Dashed lines represent fluorescent emission peaks.

**S16 Fig. Fluorescent emission spectra of Oxudercidae.** Dashed lines represent fluorescent emission peaks.

**S17 Fig. Fluorescent emission spectra of Tripterygiidae.** Dashed lines represent fluorescent emission peaks.

**S18 Fig. Fluorescent emission spectra of Blenniidae.** Dashed lines represent fluorescent emission peaks.

**S19 Fig. Fluorescent emission spectra of Antennariidae.** Dashed lines represent fluorescent emission peaks.

**S20 Fig. Fluorescent emission spectra of Cynoglossidae.** Dashed lines represent fluorescent emission peaks.

**S21 Fig. Fluorescent emission spectra of Bothidae.** Dashed lines represent fluorescent emission peaks.

**S22 Fig. Fluorescent emission spectra of Soleidae.** Dashed lines represent fluorescent emission peaks.

**S23 Fig. Fluorescent emission spectra of Synodontidae.** Dashed lines represent fluorescent emission peaks.

